# Gene signatures characterizing driver mutations in lung squamous carcinoma are predictive of the progression of pre-cancer lesions

**DOI:** 10.1101/2025.07.29.664792

**Authors:** Yupei Lin, Venugopalareddy Mekala, Jianrong Li, Xiang Wang, Muhammad Aminu, Jia Wu, Jianjun Zhang, Robert Taylor Ripley, Christopher I. Amos, Chao Cheng

## Abstract

Patients with lung squamous cell carcinoma (LUSC) are often diagnosed at advanced stages, limiting opportunities for early intervention. LUSC develops through a multistep progression from low-grade lesions to high-grade lesions, including carcinoma in situ (CIS), of which approximately half progress to invasive cancer while the other half regress. Although frequent mutations and copy number alterations have been documented in LUSC and observed in precursor lesions, their prognostic significance in precancerous stages remains largely unexplored.

In this study, we leveraged gene expression data from LUSC tumors in The Cancer Genome Atlas (TCGA) to derive transcriptional signatures corresponding to 34 key driver genomic aberrations, including mutations, amplifications, and deletions. These tumor-derived gene signatures were then applied to precancerous lesion datasets to assess their ability to characterize developmental stages and predict progression risk.

We found that many of these signatures increased progressively across lesion stages, reflecting their roles in early tumorigenesis. In particular, several signatures accurately predicted which CIS lesions would progress to invasive cancer. Furthermore, these signature scores were more strongly associated with patient prognosis in LUSC than the presence of genomic aberrations alone.

We also examined the relationship between driver-associated signatures and the tumor immune microenvironment. Signature scores were significantly correlated with immune features such as immune cell infiltration and immune checkpoint gene expression, including CD274 (PD-L1). Interestingly, these associations varied across lesion stages, indicating dynamic immune interactions during cancer evolution.

Together, our findings demonstrate that tumor-derived driver gene expression signatures provide valuable insight into the biology and progression risk of precancerous lesions, offering potential utility for early detection and intervention strategies in LUSC.

**Novelty and Impact:** In this study, we developed gene expression signatures using lung squamous cell carcinoma (LUSC) transcriptomic and genomic data and apply them to characterize precancerous lesions. These signatures showed high performance in predicting the progression risk of carcinoma in situ (CIS), offering a valuable tool for personalized prevention and early intervention in LUSC. These findings have the potential to improve early lung cancer detection and provide new insights into the tumor microenvironment in the context of precancer evolution of LUSC.

## Introduction

Lung squamous cell carcinoma (LUSC) accounts for approximately 20% of primary lung cancers in the US, making it the second most prevalent subtype, following lung adenocarcinoma.^1–3^ Compared with lung adenocarcinoma, LUSC is more aggressive with a lower median survival rate. A key factor contributing to the low survival rate is its frequent diagnosis at advanced stages.^4–8^ To improve prognosis, early detection and continuous surveillance are essential. LUSC originates from normal bronchial tissue in the central airways,^9^ and proceeds through several sequential precancer stages, including hyperplasia, metaplasia, mild dysplasia, moderate dysplasia, severe dysplasia, and carcinoma-in-situ (CIS).^8,9^ Among these, CIS represents a preinvasive stage where carcinoma cells are confined to the uppermost layer of cells in the bronchi, bronchioles, or alveoli, without penetrating deeper tissue layers.^10^ Approximately 50% of patients diagnosed with CIS progress to LUSC within three years, while 30% experience spontaneous regression.^11^ To date, there is no effective method to predict CIS progression; however, recent studies suggested that transcriptomic profiles are highly informative in this regard.^8,9,11,12^

The initiation and progression of cancer are driven by numerous genomic aberrations, including mutations and copy number changes in oncogenes and tumor suppressor genes.^13–15^ The Cancer Genome Atlas (TCGA) lung cancer group identified recurrent mutations in genes such as TP53, CDKN2A, and PIK3CA; amplifications of regions including chromosome 3q (containing SOX2 and PIK3CA), EGFR, and NFE2L2; and deletions in FOXP1, PTEN, and NF1.^5,16,17^ Interestingly, these genomic events showed limited capability in prognostic stratification of LUSC patients.^14,18^ This limitation is due to the complexity of tumorigenesis, where tumor initiation and progression are driven by specific oncogenic pathways that can be dysregulated through alternative mechanisms beyond the primary genomic aberrations. For instance, although TP53 mutations occur in over 50% of tumors, the P53 pathway can also be inactivated through TP53 gene loss, hypermethylation of the TP53 promoter, or alterations in other pathway-related genes.^15,19,20^ Previous studies have demonstrated that gene signatures representing deregulated oncogenic pathways provided more accurate prognostic indicators than mutation status themselves in multiple cancer types.^21–24^ These driver pathway-based characterizations offer superior guidance for devising targeted therapeutic interventions compared to focusing solely on genomic aberrations.^25^

It has been shown that many driver mutations occur at the early stage of cancer development.^13,26–28^ In fact, some genomic alterations in LUSC driver genes, such as PIK3CA, CDKN2A, and, SOX2 have been detected in pre-cancer lesions.^11,29–32^ Roberts et al reported that the activation of PI3K/Akt pathways occur during the transition from low- to high-grade premalignant lesions, while CDK4/cyclin-D1 indicates an early onset at the stages of metaplasia and moderate dysplasia.^8^ However, there is a lack of research on the extent to which these genomic aberrations present in precancer stages influence the progression of these lesions and how we can quantitatively characterize pathway changes during cancer progression.

In this study, we hypothesized that genomic signatures derived from tumor data can be applied to precancerous lesions to infer the progression of precancerous lesions to more advanced states including metastatic cancers. To test this hypothesis, we integrated multi-omics data from TCGA to define gene signatures for 34 genomic aberrations frequently observed in LUSC, including 10 gene mutations, 3 chromosomal region deletions, and 21 chromosomal region amplifications. When applied to multiple precancerous datasets, these signatures exhibited dynamic changes throughout distinct precancerous stages, highlighting their roles in cancer evolution. Notably, some signatures predicted the progression risk of CIS samples with high accuracy. Additionally, we observed significant correlations between these signatures and immunological features in the tumor microenvironment (TME) during cancer progression. Our study provides a valuable framework for characterizing precancerous lesions and monitoring their progression to LUSC by using gene signatures associated with driver genomic aberrations.

## Material and methods

### Datasets

All data summary is presented in Table 1. The RNA-sequencing (RNA-seq) data for 501 LUSC tumor samples from TCGA were downloaded from FireBrowse.^5,33^ This dataset provided the normalized expression values for 20,501 genes, represented as the RNA-Seq by Expectation-Maximization (RSEM) values and included associated clinical data.^34^ All the other gene expression datasets are generated by microarrays and downloaded from the Gene Expression Omnibus (GEO) public repository.^35^ GSE157011 includes gene expression profiles for 249 LUSC samples with matched overall survival and other clinical information. ^26^ The GSE108124 provide gene expression profiles for 33 CIS samples, including 17 progressive and 16 regressive samples.^11^ GSE33479 comprise gene expression profiles for 122 endobronchial biopsies at different stages including normal bronchial tissue, hyperplasia, metaplasia, mild dysplasia, moderate dysplasia, severe dysplasia, CIS, and LUSC.^7^ These data are provided as normalized expression profiles at the probe-sets level, which were converted into gene expression profiles. For genes with multiple probesets in the microarray platform, the one with maximum average hybridization signals across all samples is selected to represent gene expression.

**Table 1.** Datasets of this study.

| GEO accession | Study PMID | Sample Size | Clinical category |
| --- | --- | --- | --- |
| TCGA | 22960745 | 501 | LUSC |
| GSE157011 | 32717408 | 249 | LUSC |
| GSE33479 | 31243362 | 122 | Developmental stage |
| GSE108124 | 30664780 | 33 | CIS |

Summary of the datasets included in this study, detailing their corresponding Study ID, sample size, and sample type.

The TCGA level-3 somatic mutation data were downloaded from the TCGA data portal (http://cancergenome.nih.gov/). This cohort contains 178 samples with 21939 mutated genes from whole exome sequencing.^36^ The following categories of somatic mutations are considered to have functional effects: frame shift insertion or deletion, in frame deletion, missense mutation, splice site mutation and nonsense mutation. The level 3 LUSC copy number variation (CNV) data measured by the Affymetrix Genome-Wide Human SNP Array is used. There are 501 LUSC cases and 23311 genes in the TCGA CNV profile.

### Selected driver genomic aberrations frequently occurring in LUSC

In this study, we started with 513 cancer genes collected by The Catalogue of Somatic Mutations in Cancer (COSMIC) Cancer Gene Census (CGC).^13^ Genomic aberrations associated with these cancer genes are selected based on their frequency in the TCGA-LUSC data. Based on the processed somatic mutation data, we selected 10 cancer genes that are functionally mutated in at least 10% of LUSC samples. Similarly, we selected 46 cancer genes that are amplified (43 genes) or deleted (3 genes) in at least 10% LUSC samples based on processed CNV data. Of note, some of the frequently amplified genes are located proximally in the genome. As a result, their amplifications are mostly caused by the same set of broad chromosome band amplification events. Thus, we merged these genes to obtain independent amplification events named by their locations in the genome. Specifically, the following frequently amplified cancer gene groups were defined: chr1q21:23, chr2p15:16, chr3q12:13, chr3q21:22, chr3q25, chr3q26:27, chr5p13, chr8q22:24, chr8p11, chr12p13, (Supp. Table 1). The amplification status of the most frequently amplified genes within a group was selected to represent the amplification status of the corresponding gene group in each LUSC sample. Ultimately, we obtained 34 genomic aberrations including somatic mutations of 10 genes, deletions of 3 genes, and amplifications of genes (n=11) or gene groups (n=10).

### Define gene-signatures for driver genomic aberrations

Here we describe how we defined gene signatures for each of these 34 driver genomic aberrations based on the LUSC transcriptomic data from TCGA (Supp. Note 1). First, we performed a univariable regression model for each driver events to identify the informative genes. We encoded a binary indicator variable for each genomic feature in each sample (*X_i_* = 1: presence of mutation; *X*_*i*_ = 0: absence of mutation). From the regression summary, we obtained the coefficient matrix *β*_*i*x*m*_ and the p-values matrix *P*_*g*x*m*_ for all of g genes and m events. Informative genes are those significant entries in *P*_*g*x*m*_ matrix. Next, we defined weight profiles consisting of all genes. We constructed a pair of weighted profiles respectively for up-regulated and down-regulated genes. For a particular driver event with coefficient *β*_*k*_ and p-value *P*_*k*_, we defined the weight by the following *β_k_* >0 = *-log_l0_* (*P_k_*) and *β*_*k*_ <0= log_*l0*_ (*P*_*k*_). To avoid the extreme values, we trimmed all weights greater than 10 to 10. Please see Supp. Note 1 for method details.

### Calculate sample-specific gene signature score based on gene expression profiles

We used the previously developed computational algorithm BASE to calculate rank-based similarity score between TCGA-LUSC derived gene expression score and gene expression profile from independent validation datasets.^37^ We followed these procedures: 1) Given an expression profile, we sorted genes in descending order as e = (*e*_l_, *e*_2_,…, *e*_*n*_), n is the total number of genes. 2) We validated whether genes with high weights showed a skewed distribution and quantified the distribution difference. 3)The BASE algorithm then calculated maximum deviations against the permutation by randomly selecting samples resulting in *S*^*up*^ and *S*^*dn*^. The final gene-signature score was defined as: S = *S*^*up*^ - *S*^*dn*^ where for each gene-signature, we determined the final score with the subtraction of the enrichment score of up-regulated models and down-regulated models for g. Please see Supp. Note 1 for method details.

### Compute the infiltration level of immune cells based on gene expression profiles

Similarly, we used BASE to infer immune infiltration by comparing the transformed reference immune cell profiles with patient gene expression data as previously published.^37,38^ This application of the BASE algorithm estimated the immune cells’ infiltration levels by examining the expression levels of immune-cell-specific genes. Thus, we derived the infiltration scores of major immune cell types including memory B cells, CD4+ T cells, CD8+ T cells, naïve B cells, NK cell and macrophage. Please see Supp. Note 1 for BASE algorithm details.

### Statistical analysis

All statistical analyses were conducted in the R software. We use “coxph” function for univariable and multivariable Cox regression. R package “survival” was implemented to perform survival analyses. The “survdiff” function was used to compare patient groups using log-rank test. The “wilcox.test” and the “t.test” functions were used to compare two groups of selected measurement. False discovery rate (FDR)-adjusted p-values were calculated using the Benjamini-Hochberg method via the p.adjust() function in R. Supplementary lasso regression was performed using the “glmnet” package applying penalty term^39^. Correlation analysis utilized the function “cor.test” with default Spearman coefficients. The package “ggplot2” was used to generate boxplots and Kaplan-Meier survival time curves. The Mann-Kendall trend test was employed to analyze data collected over time for consistently increasing or decreasing trends (monotonic) in response values. Specifically, the function “MannKendall” from the R package “Kendall” was used to implement this test.^40^ Hierarchical clustering with default distance (‘Euclidean’) and method (‘complete’), and heatmap visualization for Figure 5, was performed using ComplexHeatmap. Additionally, the K-means algorithm was utilized to divide samples into immune hot and immune cold categories, with the parameter ‘column_km’ set to 2. A p-value of 0.05 serves as the significant threshold for statistical tests, unless otherwise noted.

## Results

### Overview of this study

Tumorigenesis is driven by genomic aberrations that activate oncogenic pathways or inactivate tumor suppressive pathways, often occurring at the early stages of malignancy. While extensive cancer genomic and transcriptomic data exist, precancer data remains limited. To address this gap, we developed a novel framework (Supp. Note 2). Figure 1 left panel illustrates tumor samples from TCGA-LUSC and the development of gene signatures for driver genomic aberrations, integrating genomic data (somatic mutations and copy number variations) and transcriptomic data (RNA-seq) from LUSC samples. These signatures, which represent 34 frequent genomic aberrations (including 10 gene mutations, 3 chromosome region deletions, and 21 chromosome region amplifications), recapitulate the deregulated pathways underlying the aberrations and are detailed in Supp. Table 4. The right panel shows the application of these cancer-derived signatures to precancer data, where we found that they correlated more strongly with patient prognosis in an independent LUSC dataset. To validate their utility in distinguishing precancer stages, we applied them to multiple precancer datasets, assessing their ability to predict CIS progression and characterize immunological features in the TME.

**Figure 1:**
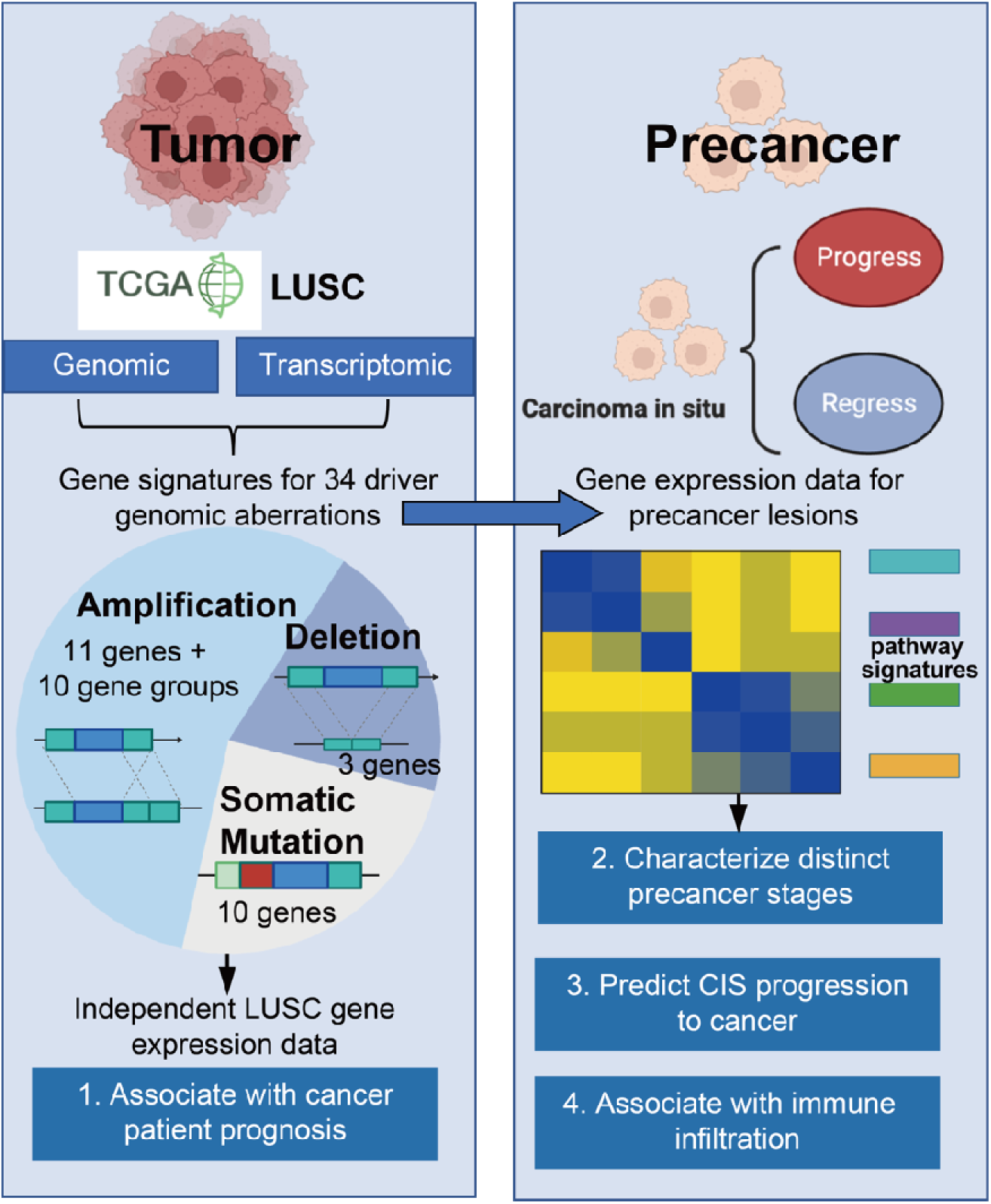
Overview of this study. Gene signatures for 34 driver genomic aberrations occurring in >10% of TCGA-LUSC samples are defined to recapitulate the underlying de-regulated oncogenic pathways. These signatures were used to independent cancer and precancer gene expression datasets to 1) examine their prognostic association, 2) characterize distinct precancer stages, 3) predict the progression of CIS to cancer, and 4) investigate the influence of driver genomic aberrations on immune infiltration.

### Prognostic association of gene signatures in lung squamous cell carcinoma

Out of the 34 LUSC genomic aberrations, two are implicated in the inactivation of CDKN2A: the mutation of CDKN2A gene and the deletion of the chromosomal region which harbors CDKN2A gene. Interestingly, the CDKN2A-MUT signature can detect both events: LUSC samples harboring mutated CDKN2A and chr9p21 deletions both showed significantly higher scores than their normal counterparts (Figure 2A). However, the chr9p21-DEL signature can only detect samples harboring chr9p2 deletions but not CDKN2A gene mutations, suggesting that it reflects the deletion effect of the whole chromosomal regions rather than CDKN2A gene alone. Similarly, the PIK3CA-MUT signature can distinguish LUSC samples with mutated PIK3CA or amplified Chr3q26:27 regions (harboring PIK3CA gene) from their normal counterparts (Figure 2B).

**Figure 2:**
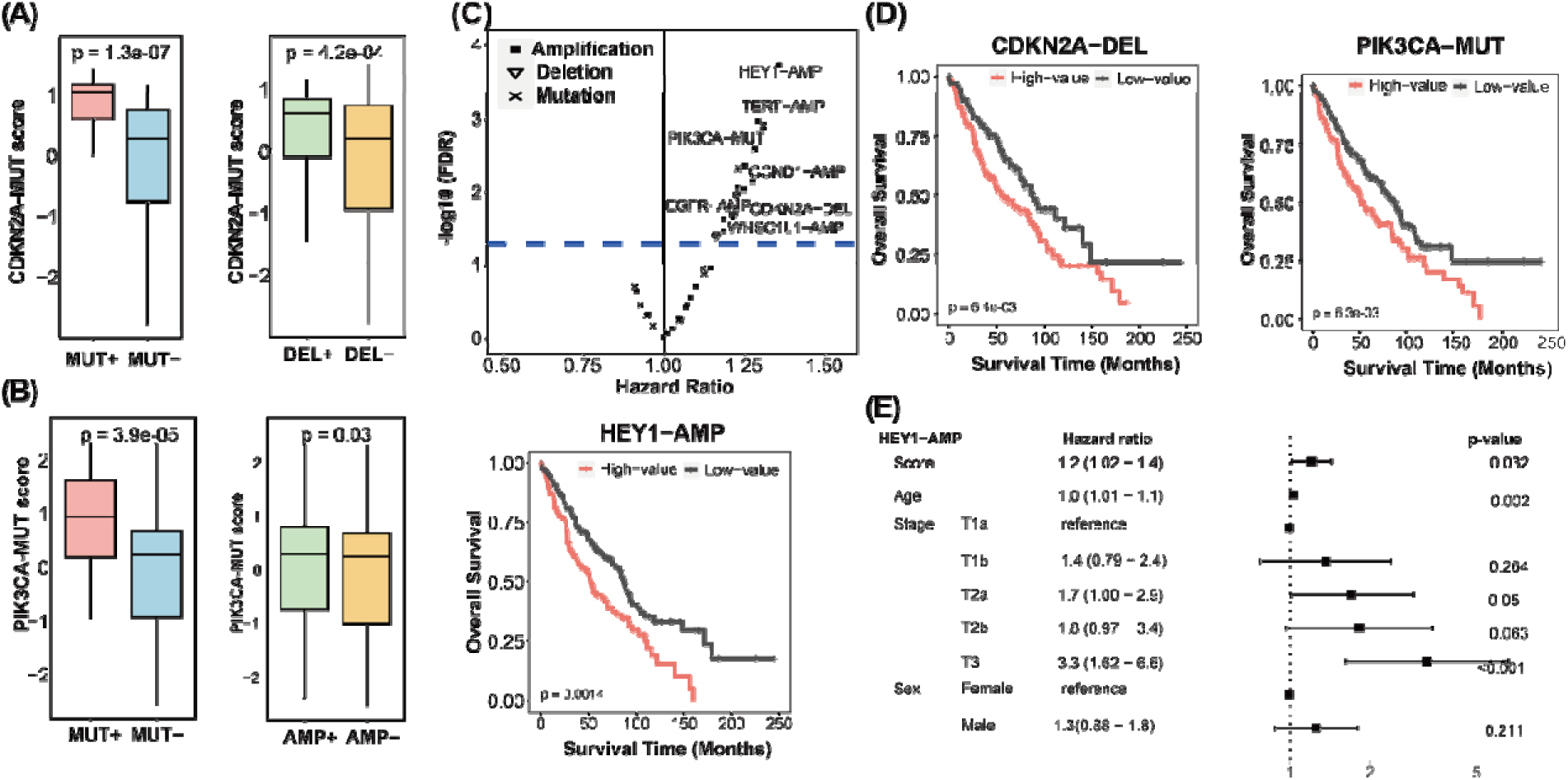
Gene signatures for driver genomic aberrations are predictive of patient prognosis in lung squamous cell carcinoma. (A) In TCGA-LUSC data, the CDKN2A mutation gene signature can not only distinguish LUSC samples with mutated CDKN2A gene (MUT+) from those without (MUT-, but also can distinguish samples with CDKN2A deletion (DEL+) from those without (DEL-). (B) Similarly, the PIK3CA-MUT gene signature can distinguish LUSC samples with mutated PIK3CA genes (MUT+) from those without (MUT-), and PIK3CA amplification (AMP+) from those without (AMP-). Note: Generally, mutations in CDKN2A gene result in loss of function, whereas PIK3CA mutations lead to gain of function; Deletions of CDKN2A and amplifications of PIK3CA commonly arise due to the loss of chr3q26 and gain of chr3q26, respectively. (C) In GSE157011-LUSC data, volcano plot indicating the gene signatures for many driver genomic aberrations are significantly associated with patient prognosis. Horizontal line indicates the significance threshold FDR<0.05. (D) Kaplan-Meier curves showing 3 example signatures (HEY1-AMP, CDKN2A-DEL and PIK3CA-MUT) can stratify patients (median signature score as the cut-off value) into subgroups with significant prognosis. (E) Forest plot indicating the HEY1-AMP signature score remains prognostically significant after adjusting for age, stage, and sex in the GSE157011-LUSC data.

We then investigated the prognostic relevance of these gene signatures in LUSC. We applied our gene signatures to an independent LUSC dataset generated from GSE15701. Univariable Cox regression analysis identified 14 signatures significantly associated with prognosis (Figure 2c, Supp. Table 2). Interestingly, all these significant signatures were negatively correlated with patient survival (Hazard ratio HR>1), indicating that the presence of these aberrations, or more accurately, the deregulation of the related pathways, was detrimental to patient prognosis. For examples, when stratified by the scores of HEY1-AMP (chr8q21), CDKN2A-DEL (chr9p21), or PIK3CA-MUT signatures (using median as the cut-off), patients with higher scores had significantly shorter overall survival time than those with lower scores (Figure 2d). Multivariable Cox regression analysis indicated that some of these signatures remained significant after adjusting for major clinical variables including age, sex, and tumor stage (Figure 2e).

### Apply gene signatures to characterize precancer samples at different stages

To validate the effectiveness of our gene signatures in precancerous samples, we first applied them to the CIS dataset GSE108124, which includes transcriptomic profiles for 33 CIS samples and their TP53 mutation status. The TP53-MUT signature showed significantly higher scores in the six CIS samples with mutated TP53 compared to those without (Figure 3a). Interestingly, for the samples without TP53 mutation (TP53WT), TP53-MUT signature provided additional stratification by significantly differentiating patients based on their eventual progression status. Next, we applied all gene signatures to another precancer dataset (GSE33479), which provided expression profiles from six different precancer stages, as well as from normal lung and LUSC samples (Figure 3b). Trend analysis revealed that 23 out of the 34 signatures exhibited a significantly monotonic trend during the evolution from early to advanced stages (P < 0.05, Mann-Kendall Trend test).

**Figure 3:**
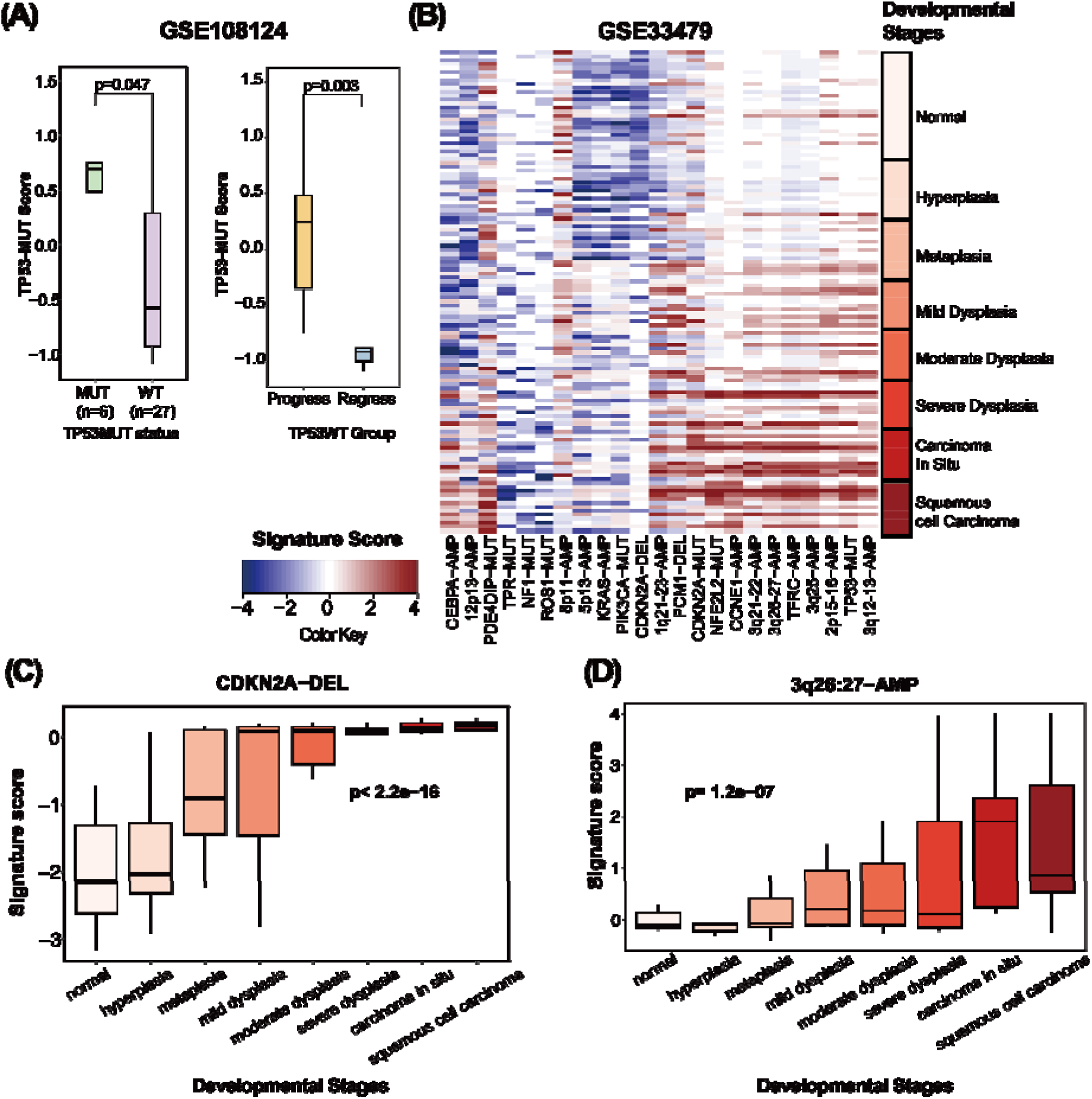
Gene signatures defined based on LUSC data can be applied to precancer gene expression data to characterize the relevant pathway activity changes across different precancer stages. (A) In GSE108124 precancer data, boxplot showing that the TP53-MUT gene signature can distinguish precancer samples with mutated TP53 gene from those without; In patients without TP53 mutation (TP53WT), TP53-MUT signature can further differentiate precancer samples on their eventual progression status. (B) In GSE33479, heatmap showing the scores of the 23 significant signatures in samples from 8 developmental precancer stages, depicted in the right color bar. Signature scores are scaled. (C) The CDKN2A-DEL gene signature scores exhibit a progressive increase across histologic stages, rising from low levels in normal and hyperplastic tissues to markedly higher levels at the metaplastic stage, suggesting a role in early cancer progression. (D) The 3q26:27-AMP gene signature scores also follow a stepwise upward trend across precancer stages, indicating potential involvement in later stages of progression toward LUSC.

Most of these signatures showed a continuous increase during precancer progression (Figure 3b). For example, the CDKN2A-DEL signature had relatively low scores in normal lung and hyperplasia samples but increased significantly at the metaplasia (P < 4.2e-04, Wilcox test compared to hyperplasia) and mild dysplasia (P < 2.7e-03, Wilcox test compared to metaplasia) stages. After these stages, the CDKN2A-DEL signature maintained high scores in subsequent stages. Another example is the chr3q26:27-AMP signature, which showed relatively low scores until the severe dysplasia stage, after which it significantly increased in CIS samples (Figure 3c). Altogether, these results suggest that gene signatures derived from LUSC samples can effectively characterize precancerous samples, providing insights into the specific stages at which genomic aberration-related pathways become active. We examined how the CDKN2A-DEL signature in TCGA relates to tumor phenotypes using molecular features from Thorsson et al. and found that higher CDKN2A-deletion scores were associated with increased proliferation (R = 0.53) and decreased macrophage regulation (R = –0.76) and lymphocyte infiltration (R = –0.63). These findings are consistent with the role of CDKN2A as a tumor suppressor that limits cell cycle progression, where its loss may drive unchecked proliferation. The negative correlations with immune-related features suggest that CDKN2A deletion may also contribute to a less immune-infiltrated, potentially immune-evasive tumor microenvironment.

### Apply gene signatures to predict the progression of lung carcinoma in situ

Having found that some driver genomic signatures were monotonically increasing during progression from the early to later precursor stages, we next investigated whether the variations of these signature scores within CIS samples inform their progression risk into cancers. First, we performed hierarchical clustering analysis using the signature scores across all CIS samples in GSE108124. As shown in Figure 4a, the signature scores separated the samples into two groups, one enriched for progressive while the other enriched for regressive samples. This indicated that many of these signatures were informative for distinguishing progressive from regressive samples. Taking the CDKN2A-MUT model as an example, we found that the samples which eventually progressed to LUSC exhibited higher scores than those that regressed (p = 7.9e-09). Subsequent waterfall plots visually depicted the score distribution for all CIS samples, revealing higher scores for progressive samples across most signatures (Figure 4c). However, some pathway signatures, like NF1-MUT, portrayed an opposite trend, with higher scores in regressive patients and lower scores in progressive patients (Figure 4d).

**Figure 4:**
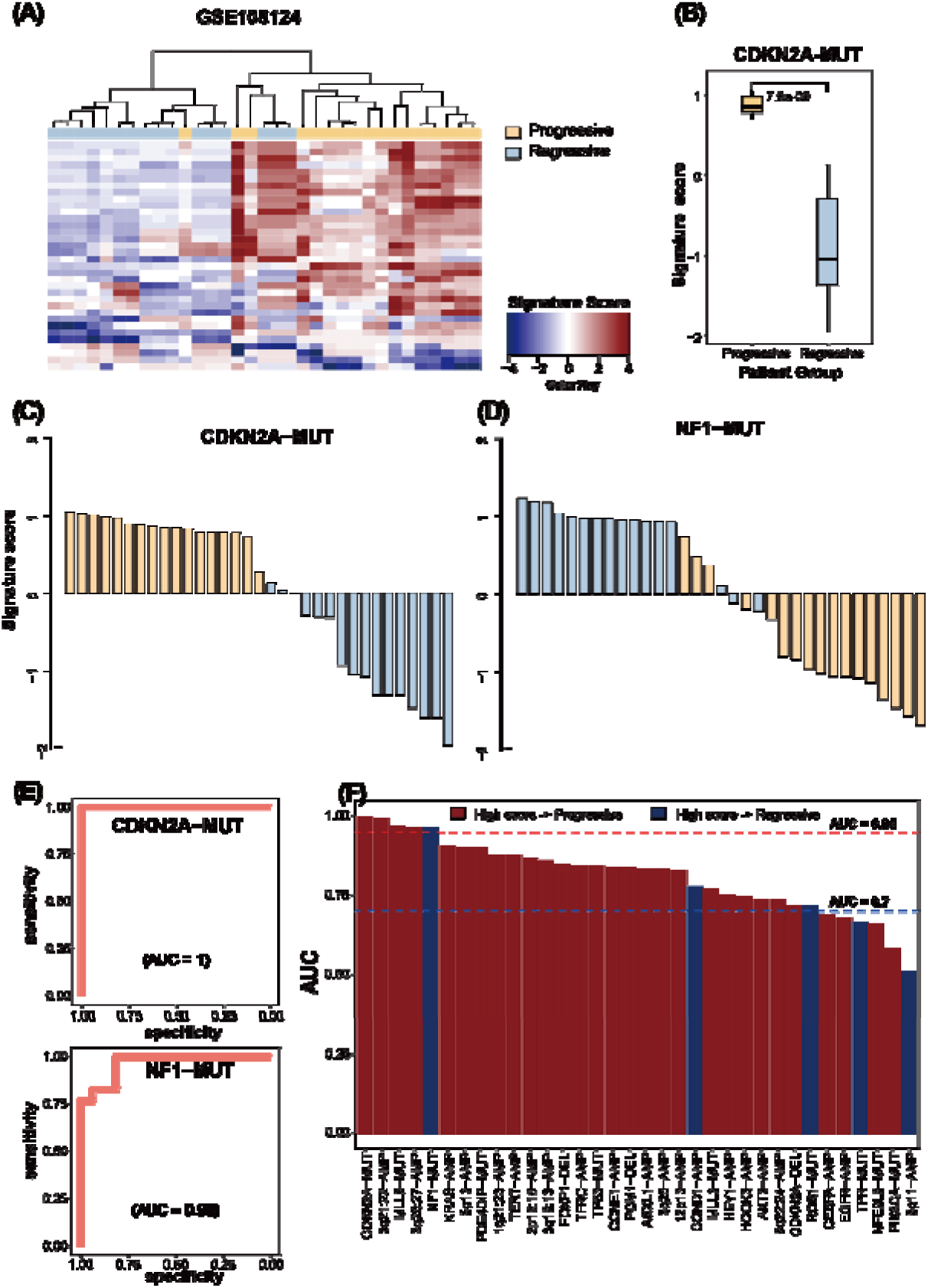
Driver gene signatures are predictive of the progression of patients with lung carcinoma in situ. (A) In GSE108124 precancer data, heatmap showing the scores of 34 signatures in carcinoma in situ samples, with progression status color-coded on the top bar. (B) Boxplot showing that progressive samples tend to have a higher CDKN2A-MUT score than regressive samples. (C) and (D) Waterfall plots displaying all CIS samples’ signature scores for CDKN2A-MUT and NF1-M T; (E) ROC plots showing that CDKN2A-MUT reaches an AUC score of 1, while NF1-MUT has an AUC of 0.96. (F) Barplots presenting AUC scores for each of our 34 gene signatures in descending order. Bars in red indicate that progressive patients tend to have higher signature scores, whereas bars in blue indicate that regressive patients tend to have higher signature scores. The blue dashed line represents the cutoff of an AUC of 0.7, and the red dashed line represents the cutoff of an AUC of 0.95.

The predictive powers of the CDKN2A-MUT and the NF1-MUT could also be visualized by their ROC curves (Figure 4e). The Area of Under the ROC curves (AUC) provided a quantification of their ability in predicting CIS progression. As shown, the two signatures respectively achieved an AUC of 1.0 and 0.96, suggesting their high prediction accuracy. Indeed, we calculated the AUC values for all driver genomic signatures and found that many of them exhibited fairly good prediction power (AUC>0.7). Out of them, 5 gave rise to an AUC greater than 0.95, including NF1-MUT, CCND1-AMP, ROS1-MUT, TPR-MUT, 8p11-AMP (Figure 4f). Most of these signatures were positively correlated with the progression with higher scores indicating higher progression risk, while 5 were negatively correlated with progression with higher scores indicating high chance of regression. These results suggested the activity of the pathways underlying these driver genomic aberrations may play critical roles in determining the progression or regression of CIS samples.

### Associate gene signatures with immunological features in the TME of precancer samples

Previous studies have shown the immune microenvironment plays a significant role in the initiation and progression of cancer.^1,41–43^ Thus, we investigated the influence of distinct driver genomic mutations on the immune microenvironment during the progression of cancer. First, we performed hierarchical clustering analysis based on the expression of immune cell marker genes in the GSE33479 multi-precursor data, which distinguished all precancerous samples into two distinct groups characterized by high (immune-hot) and low (immune-cold) immune gene expression, respectively (Figure 5a). Subsequent comparisons of the two groups revealed that most gene signatures had elevated scores in the ‘cold’ group compared to the ‘hot’ group (Figure 5b-d). This suggested that the associated driver genomic aberrations may facilitate the development of a ‘cold’ immune microenvironment, leading to the progression of precancer lesions via immune evasion. In TCGA-LUSC dataset, we observed that CDKN2A deletion and co-deletion of CDKN2A and JAK2 were significantly enriched in immune-cold tumors, with co-loss observed in 8.96% of cold samples compared to 4.84% of hot samples. These results indicate a potential synergistic effect of CDKN2A and JAK2 co-loss in shaping an immunosuppressive microenvironment. (Supp. Figure 1)

Following that, we investigated the correlation between signature scores and immune cell infiltration at different precancer stages. To this end, we deconvoluted the gene expression of all precancer samples to calculate the infiltration level of 6 major immune cell types including CD4+ T cells, Naïve B cells, memory B cells, CD8+ T cells, NK cells, and macrophages (Figure 5e-f). We focused on their correlation with driver gene signatures that are frequently studied in cancer progression, including CDKN2A-MUT, TP53-MUT, PIK3CA-MUT, chr3q26:27-AMP, KRAS-AMP, and CDKN2A-DEL. Our analysis revealed significantly negative correlations of these signatures with the infiltration level of CD8+ T cells, suggesting the presence of these driver aberrations or their equivalent is linked with a reduced CD8+ T cell infiltration across all precancer stages (Figure 5f). In contrast, we observed significantly positive correlation of these signatures with the infiltration level of macrophages in mild dysplasia, carcinoma in situ, and squamous cell carcinoma but significantly negative correlation in metaplasia samples. Such a transition of correlations might be caused by the functional alterations of macrophages from anti-tumoral M1 cells to pro-tumoral M2 cells during the evolution of cancer through different precancer stages.^6,41^

Lastly, we examined the association between these signatures and the expression of major immune regulatory genes at different precancer stages. We thoroughly calculated the spearman correlation between signatures and immune regulatory genes (Supp. Table 3). Specifically, the expression of CD274 gene (encoding programmed cell death ligand 1, PD-L1) was negatively correlated with these signatures in pre-cancerous lesions (Figure 5g). For example, we included above driver gene signatures with addition of EGFR-AMP and NFE2L2-MUT all present negative correlation at stages of hyperplasia, metaplasia, mild dysplasia, carcinoma in situ. In contrast, high PD-L1 expression tended to correspond with higher signature scores at the LUSC stage. This shift potentially reflects a less immunosuppressive microenvironment during early progression, while the reversal at the invasive stage suggests that CDKN2A loss may contribute to immune evasion through the upregulation of PD-L1. In conclusion, our signatures emphasized the complex interplay between underlying cancer pathways and immune dynamics, bridging the gap between tumor progression and the immune microenvironment.

**Figure 5:**
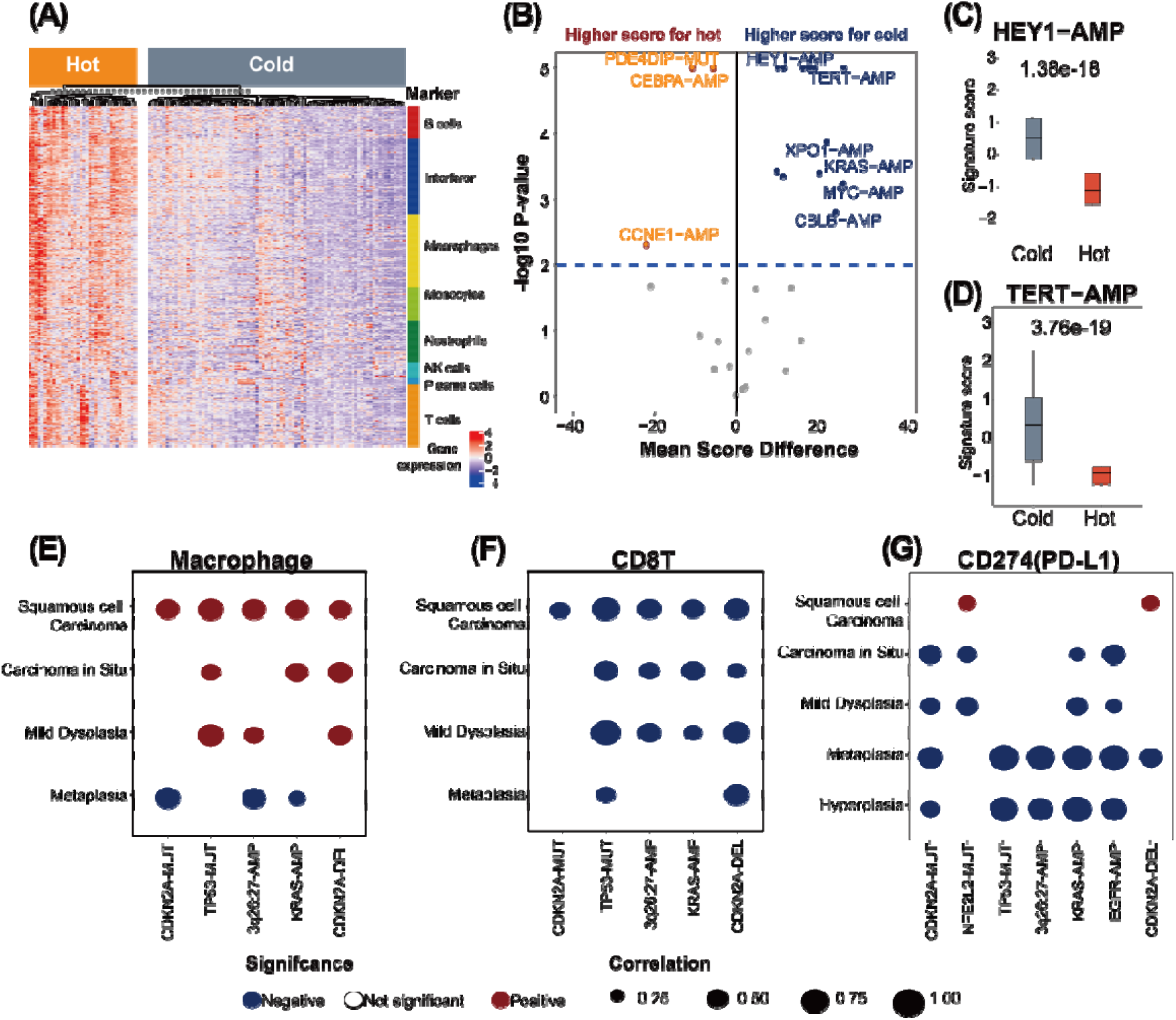
Association between gene signature and immune cells show pattern along cancer progression. (A) In GSE33479 data, heatmap showing samples categorized into the immune hot nd immune cold groups, with hierarchical clustering of marker gene expression including B cells, interferon, macrophages, monocytes, NK cells, plasma cells, and T-cells. The rightmost color bar represents the groups for each marker gene group. (B) Volcano plot illustrating significant differences in immune hot and immune cold samples. (C) and (D) Boxplots displaying HEY1-AMP and TERT-AMP, where immune cold samples tend to have a higher signature score than immune hot samples; Dot plots showing the correlation pattern between selected gene signatures and developmental stages in (E) Macrophage infiltration, (F) CD8T infiltration, and (G) PD-L1 expression (CD274). Dot size is scaled with the correlation coefficient, and color represents the directionality of correlation. Dots with a non-significant p-value are left blank.

## Discussion

The tumor-derived signatures effectively characterized and differentiate samples in pre-cancerous stages, implying that pathways associated with known driver events potentially involved early in cancer initiation. The dysregulation of cancer-related pathways is observed in both the initiation and progression of LUSC, suggesting a potential role in shaping the precancerous landscape. It is evident that in precancerous lesions, the expression profile is not predominant by a single mechanism, but rather multiple pathway activities contribute to the ultimate result.

The majority of signatures exhibited a pattern where higher scores were associated with worse prognosis in LUSC or higher progression risk in CIS. However, some signatures, such as NF1-MUT, CCND1-AMP, and ROS1-MUT, showed an opposite trend, with lower NF1-MUT scores correlating with better prognosis (Figure 4f). This suggests that certain genomic events may act as secondary mutations and interact with other pathways to produce a more favorable effect. NF1, a tumor suppressor gene that regulates Ras signaling, displays highly variable mutations across the genome.^44^ Studies have shown that while one-third of samples harbor both NF1 and TP53 mutations, NF1 mutation coexists with KRAS mutations less frequently. Interestingly, patients with only an NF1 mutation tend to have longer survival than those with concurrent NF1 and KRAS mutations.^45^ Furthermore, NF1 mutations appear to prolong patient survival in EGFR mutant/TP53 wild-type patients compared to those with EGFR/TP53 co mutations.^46^ Additional research has reported that low levels of NF1 expression are associated with resistance to EGFR-targeted therapies in lung cancer patients, suggesting a complex role of NF1 in tumorigenesis and therapy resistance.^5,47^

CDKN2A encodes p14^ARF and p16^INK4a, which regulate apoptosis and cell cycle progression.^48^ Given that anti-PD-1/PD-L1 therapies rely on intact apoptotic pathways, CDKN2A loss, particularly affecting p14^ARF which may impair response to immunotherapy.^49,50^ While prior studies have suggested that 9p21.3 deletions may reduce PD-L1 expression via broader 9p loss extending into 9p24.1,^51^ our CDKN2A-DEL signature primarily reflects the dysregulated pathways associated with CDKN2A loss (Supp. Table 7). Nevertheless, we acknowledge that most CDKN2A losses result from the deletion of a larger chromosomal segment encompassing many other genes. Therefore, the CDKN2A-DEL signature may also capture effects mediated by the loss of these neighboring genes. Despite the limited number of samples with stringent Chr9p21 loss, we observed a strong concordance between the CDKN2A-DEL and Chr9p21-DEL signatures. Most Chr9p21-deleted samples also harbored CDKN2A deletion, and the two signatures showed high correlation in gene weights and score patterns across datasets. Their associations with shorter survival and increased progression risk were also similar. These results suggest that the CDKN2A-DEL signature may reflect broader deletion effects across the Chr9p21 region. However, our framework does not fully separate the independent effects of CDKN2A loss from nearby genes in the Chr9p21-24 region and current study still focused on the gene-level CDKN2A-DEL signature. Furthermore, we found that the CDKN2A-DEL signature correlated with deletion segment size across the 9p21 region (R = 0.47).

Although our study does not include direct mechanistic validation for CDKN2A deletion pathway during the pre-cancer stages, the observed increase in CDKN2A-DEL signature scores rising from low levels in lower grades to markedly higher levels at the metaplastic stage suggests a biological transition toward increased proliferation and immune evasion associated with CDKN2A loss. These findings potentially suggest a functional role for CDKN2A loss in driving histologic progression and immune modulation.

The intricate relationships in TME play a crucial role in the lung cancer progression leading to various methods for stratifying levels of immunogenicity. Immune “hot” regions typically express high levels of cytotoxic lymphocytes and immune activation markers, while “cold” immune samples tend to have worse survival outcome.^52,53^ Parra et al. demonstrated in LUSC, a cold tumor microenvironment is usually correlated with a high burden of chromosomal copy-number variants and is associated with a reduced benefit from immune checkpoint inhibitors.^54^ In our study, we found that most samples exhibited high risk score on cold immune environment, while a few signatures, such as PDE4DIP-MUT, had higher signature scores in immune hot samples. Furthermore, we observed enrichment of CDKN2A deletion and CDKN2A/JAK2 co-loss in immune cold LUSC, supporting its role in shaping an immunosuppressive microenvironment, as also noted in other studies.^55^ These findings suggest a context-dependent role of 9p loss in immune evasion, particularly during the transition from precancer to invasive disease.

In GSE108124, gene expression profiles were generated intentionally for index CIS lesion biopsies, representing the latest biopsy that were collected preceding progress to invasive cancer or regress to low-grade dysplasia or normal epithelium.^11^ Compare with biopsies collected at other times, the molecular profiling of these index CIS samples are more informative for predicting the progression risk of CIS patients. This may explain the overall high accuracy of our driver genomic signatures for predicting CIS progression in the data. In the future it would be interesting to perform survival analysis to associate signature scores with progression-free survival of CIS samples.

The gene signatures identified in this study offer promising clinical utility for stratifying patients with precancerous lesions based on their risk of progression to LUSC. As CIS lesions are commonly managed with surgical resection, integrating our predictive model into clinical workflows could enhance postoperative risk assessment. Patients with high-risk CIS should adopt more intensive surveillance strategies after surgery to prevent recurrence or progression. However, results from this study are based on retrospective data without experimental validation. A more careful validation using prospective data from independent patient cohorts are needed to translate these finding into clinical applications.

In summary, we conducted a comprehensive study utilizing a set of 34 driver-aberration signatures derived from TCGA-LUSC data. We validated the prognostic relevance of these driver genomic signatures and demonstrated its potential utility in characterizing precancer samples of LUSC. These signatures are predictive of the progression risk of CIS samples, and therefore has the potential to aid clinicians in making decisions for personalized CIS surveillance and treatment.

## Abbreviations

CIS: Carcinoma-in-situ
CNV: Copy Number Variation
COSMIC-CGC: The Catalogue of Somatic Mutations Cancer Gene Census
GEO: Gene Expression Omnibus
LUSC: Lung Squamous Cell Carcinoma
NK: Natural Killer cells
TCGA: The Cancer Genome Atlas
TME: Tumor Microenvironment.

## Acknowledgements

This study was supported by the Cancer Prevention Research Institute of Texas (Cheng: RR180061 and Amos: RR170048) and the National Cancer Institute of the National Institutes of Health (Cheng: 1R01CA269764). CC is a CPRIT Scholar in Cancer Research. This work was supported by Integrative Analysis of Lung cancer Etiology and Risk (Amos: U19CA203654). This grant was supported by an CDC U24 (Ripley: 2U24OH009077-15-00). National Cancer Institute of the National Institute of Health Research Project Grant (R01CA234629), the AACR-Johnson & Johnson Lung Cancer Innovation Science Grant (18-90-52-ZHAN), the MD Anderson Physician Scientist Program, MD Anderson Lung Cancer Moon Shot Program. All figures included in this manuscript are created by authors of this paper. Adobe Illustrator is used for final labeling. Fig. 1 is created with BioRender.

## Data Availability Statement

All data can be publicly downloaded based on the GEO accession ID provided in the materials and methods. All computational codes for this study can be accessed from <u>github.com/YPeiLin/Carcinoma-in-situ-signature</u>. Further information is available from the corresponding author upon request.

## Authors’ contributions

Y.L.: Methodology, Software, Validation, Formal analysis, Writing - Original Draft, Visualization; V.M, J.L, X.W.: Visualization, Writing –Review & Editing; M.A, J.W, J.Z, R.R: Resources, Writing –Review & Editing; C.A,C.C: Conceptualization, Methodology, Investigation, Writing - Review & Editing, Project administration.

## Ethics Statement

All data can be publicly downloaded based on GEO accession ID, no additional ethical approval was required for this study. The data were analyzed following the relevant guidelines and regulations.

## References

1. Lau SCM, Pan Y, Velcheti V, Wong KK. Squamous cell lung cancer: Current landscape and future therapeutic options. Cancer Cell 2022;40:1279–93.

2. Barta JA, Powell CA, Wisnivesky JP. Global Epidemiology of Lung Cancer. Ann Glob Health 2019;85.

3. Drilon A, Rekhtman N, Ladanyi M, Paik P. Squamous-cell carcinomas of the lung: emerging biology, controversies, and the promise of targeted therapy. Lancet Oncol 2012;13:e418–26.

4. Cancer Genome Atlas Research Network. Comprehensive molecular profiling of lung adenocarcinoma. Nature 2014;511:543–50.

5. Cancer Genome Atlas Research Network. Comprehensive genomic characterization of squamous cell lung cancers. Nature 2012;489:519–25.

6. Liu J, Lichtenberg T, Hoadley KA, Poisson LM, Lazar AJ, Cherniack AD, Kovatich AJ, Benz CC, Levine DA, Lee A V, Omberg L, Wolf DM, et al. An Integrated TCGA Pan-Cancer Clinical Data Resource to Drive High-Quality Survival Outcome Analytics. Cell 2018;173:400–416.e11.

7. Mascaux C, Angelova M, Vasaturo A, Beane J, Hijazi K, Anthoine G, Buttard B, Rothe F, Willard-Gallo K, Haller A, Ninane V, Burny A, et al. Immune evasion before tumour invasion in early lung squamous carcinogenesis. Nature 2019;571:570–5.

8. Roberts M, Ogden J, Hossain ASM, Chaturvedi A, Kerr ARW, Dive C, Beane JE, Lopez-Garcia C. Interrogating the precancerous evolution of pathway dysfunction in lung squamous cell carcinoma using XTABLE. Elife 2023;12.

9. Pennycuick A, Teixeira VH, AbdulJabbar K, Raza SEA, Lund T, Akarca AU, Rosenthal R, Kalinke L, Chandrasekharan DP, Pipinikas CP, Lee-Six H, Hynds RE, et al. Immune Surveillance in Clinical Regression of Preinvasive Squamous Cell Lung Cancer. Cancer Discov 2020;10:1489–99.

10. Gupta A, Harris K, Dhillon SS. Role of bronchoscopy in management of central squamous cell lung carcinoma in situ. Ann Transl Med 2019;7:354.

11. Teixeira VH, Pipinikas CP, Pennycuick A, Lee-Six H, Chandrasekharan D, Beane J, Morris TJ, Karpathakis A, Feber A, Breeze CE, Ntolios P, Hynds RE, et al. Deciphering the genomic, epigenomic, and transcriptomic landscapes of pre-invasive lung cancer lesions. Nat Med 2019;25:517–25.

12. Lin Y, Burt BM, Lee H-S, Nguyen TT, Jang H-J, Lee C, Hong W, Ripley RT, Amos CI, Cheng C. Clonal gene signatures predict prognosis in mesothelioma and lung adenocarcinoma. NPJ Precis Oncol 2024;8:47.

13. Sondka Z, Bamford S, Cole CG, Ward SA, Dunham I, Forbes SA. The COSMIC Cancer Gene Census: describing genetic dysfunction across all human cancers. Nat Rev Cancer 2018;18:696– 705.

14. Zhao Y, Varn FS, Cai G, Xiao F, Amos CI, Cheng C. A P53-Deficiency Gene Signature Predicts Recurrence Risk of Patients with Early-Stage Lung Adenocarcinoma. Cancer Epidemiol Biomarkers Prev 2018;27:86–95.

15. Joerger AC, Fersht AR. The p53 Pathway: Origins, Inactivation in Cancer, and Emerging Therapeutic Approaches. Annu Rev Biochem 2016;85:375–404.

16. Yoon HI, Park KH, Lee E-J, Keum KC, Lee CG, Kim CH, Kim YB. Overexpression of SOX2 Is Associated with Better Overall Survival in Squamous Cell Lung Cancer Patients Treated with Adjuvant Radiotherapy. Cancer Res Treat 2016;48:473–82.

17. Jeon T, Oh UJ, Min J, Kim C. Gene-level dissection of chromosome 3q locus amplification in squamous cell carcinoma of the lung using the nCounter assay. Thorac Cancer 2023;14:2635– 41.

18. Cheng C, Zhao Y, Schaafsma E, Weng Y-L, Amos C. An EGFR signature predicts cell line and patient sensitivity to multiple tyrosine kinase inhibitors. Int J Cancer 2020;147:2621–33.

19. Kruse J-P, Gu W. Modes of p53 regulation. Cell 2009;137:609–22.

20. Rivlin N, Brosh R, Oren M, Rotter V. Mutations in the p53 Tumor Suppressor Gene: Important Milestones at the Various Steps of Tumorigenesis. Genes Cancer 2011;2:466–74.

21. Zolotovskaia MA, Tkachev VS, Seryakov AP, Kuzmin D V, Kamashev DE, Sorokin MI, Roumiantsev SA, Buzdin AA. Mutation Enrichment and Transcriptomic Activation Signatures of 419 Molecular Pathways in Cancer. Cancers (Basel) 2020;12.

22. Du K, Wei S, Wei Z, Frederick DT, Miao B, Moll T, Tian T, Sugarman E, Gabrilovich DI, Sullivan RJ, Liu L, Flaherty KT, et al. Pathway signatures derived from on-treatment tumor specimens predict response to anti-PD1 blockade in metastatic melanoma. Nat Commun 2021;12:6023.

23. Rogers SJ, Box C, Harrington KJ, Nutting C, Rhys-Evans P, Eccles SA. The phosphoinositide 3-kinase signalling pathway as a therapeutic target in squamous cell carcinoma of the head and neck. Expert Opin Ther Targets 2005;9:769–90.

24. Schwaederle M, Elkin SK, Tomson BN, Carter JL, Kurzrock R. Squamousness: Next-generation sequencing reveals shared molecular features across squamous tumor types. Cell Cycle 2015;14:2355–61.

25. Le Tourneau C, Delord J-P, Gonçalves A, Gavoille C, Dubot C, Isambert N, Campone M, Trédan O, Massiani M-A, Mauborgne C, Armanet S, Servant N, et al. Molecularly targeted therapy based on tumour molecular profiling versus conventional therapy for advanced cancer (SHIVA): a multicentre, open-label, proof-of-concept, randomised, controlled phase 2 trial. Lancet Oncol 2015;16:1324–34.

26. Bueno R, Richards WG, Harpole DH, Ballman K V, Tsao M-S, Chen Z, Wang X, Chen G, Chirieac LR, Chui MH, Franklin WA, Giordano TJ, et al. Multi-Institutional Prospective Validation of Prognostic mRNA Signatures in Early Stage Squamous Lung Cancer (Alliance). J Thorac Oncol 2020;15:1748–57.

27. Laville D, Casteillo F, Yvorel V, Tiffet O, Vergnon J-M, Péoc’h M, Forest F. Immune Escape Is an Early Event in Pre-Invasive Lesions of Lung Squamous Cell Carcinoma. Diagnostics (Basel) 2020;10.

28. Rosenthal R, Cadieux EL, Salgado R, Bakir M Al, Moore DA, Hiley CT, Lund T, Tanić M, Reading JL, Joshi K, Henry JY, Ghorani E, et al. Neoantigen-directed immune escape in lung cancer evolution. Nature 2019;567:479–85.

29. Rooney M, Devarakonda S, Govindan R. Genomics of squamous cell lung cancer. Oncologist 2013;18:707–16.

30. Liu Y, Yin N, Wang X, Khoor A, Sambandam V, Ghosh AB, Fields ZA, Murray NR, Justilien V, Fields AP. Chromosome 3q26 Gain Is an Early Event Driving Coordinated Overexpression of the PRKCI, SOX2, and ECT2 Oncogenes in Lung Squamous Cell Carcinoma. Cell Rep 2020;30:771–782.e6.

31. Balsara BR, Testa JR. Chromosomal imbalances in human lung cancer. Oncogene 2002;21:6877–83.

32. Yamamoto H, Shigematsu H, Nomura M, Lockwood WW, Sato M, Okumura N, Soh J, Suzuki M, Wistuba II, Fong KM, Lee H, Toyooka S, et al. PIK3CA mutations and copy number gains in human lung cancers. Cancer Res 2008;68:6913–21.

33. http://firebrowse.org/. Firebrowse: Broad Institute TCGA Genome Data Analysis Center. 2016;

34. Li B, Dewey CN. RSEM: accurate transcript quantification from RNA-Seq data with or without a reference genome. BMC Bioinformatics 2011;12:323.

35. Edgar R, Domrachev M, Lash AE. Gene Expression Omnibus: NCBI gene expression and hybridization array data repository. Nucleic Acids Res 2002;30:207–10.

36. Cancer Genome Atlas Research Network. Comprehensive molecular profiling of lung adenocarcinoma. Nature 2014;511:543–50.

37. Cheng C, Yan X, Sun F, Li LM. Inferring activity changes of transcription factors by binding association with sorted expression profiles. BMC Bioinformatics 2007;8:452.

38. Varn FS, Tafe LJ, Amos CI, Cheng C. Computational immune profiling in lung adenocarcinoma reveals reproducible prognostic associations with implications for immunotherapy. Oncoimmunology 2018;7.

39. Friedman J, Hastie T, Tibshirani R. Regularization Paths for Generalized Linear Models via Coordinate Descent. J Stat Softw 2010;33:1–22.

40. Kendall: Kendall rank correlation and Mann-Kendall trend test. . 2023;

41. Prado-Garcia H, Romero-Garcia S, Aguilar-Cazares D, Meneses-Flores M, Lopez-Gonzalez JS. Tumor-induced CD8+ T-cell dysfunction in lung cancer patients. Clin Dev Immunol 2012;2012:741741.

42. AbdulJabbar K, Raza SEA, Rosenthal R, Jamal-Hanjani M, Veeriah S, Akarca A, Lund T, Moore DA, Salgado R, Al Bakir M, Zapata L, Hiley CT, et al. Geospatial immune variability illuminates differential evolution of lung adenocarcinoma. Nat Med 2020;26:1054–62.

43. Zheng X, Weigert A, Reu S, Guenther S, Mansouri S, Bassaly B, Gattenlöhner S, Grimminger F, Pullamsetti S, Seeger W, Winter H, Savai R. Spatial Density and Distribution of Tumor-Associated Macrophages Predict Survival in Non-Small Cell Lung Carcinoma. Cancer Res 2020;80:4414–25.

44. Redig AJ, Capelletti M, Dahlberg SE, Sholl LM, Mach S, Fontes C, Shi Y, Chalasani P, Jänne PA. Clinical and Molecular Characteristics of NF1-Mutant Lung Cancer. Clin Cancer Res 2016;22:3148–56.

45. Tlemsani C, Pécuchet N, Gruber A, Laurendeau I, Danel C, Riquet M, Le Pimpec-Barthes F, Fabre E, Mansuet-Lupo A, Damotte D, Alifano M, Luscan A, et al. NF1 mutations identify molecular and clinical subtypes of lung adenocarcinomas. Cancer Med 2019;8:4330–7.

46. Tian H-X, Chen Z-H, Jie G-L, Wang Z, Yan H-H, Wu S-P, Zhang S-L, Lu D-X, Zhang X-C, Wu Y-L. Prognostic features and comprehensive genomic analysis of NF1 mutations in EGFR mutant lung cancer patients. Cancer Med 2023;12:396–406.

47. de Bruin EC, Cowell C, Warne PH, Jiang M, Saunders RE, Melnick MA, Gettinger S, Walther Z, Wurtz A, Heynen GJ, Heideman DAM, Gómez-Román J, et al. Reduced NF1 expression confers resistance to EGFR inhibition in lung cancer. Cancer Discov 2014;4:606–19.

48. Zhao R, Choi BY, Lee M-H, Bode AM, Dong Z. Implications of Genetic and Epigenetic Alterations of CDKN2A (p16(INK4a)) in Cancer. EBioMedicine 2016;8:30–9.

49. Gutiontov SI, Turchan WT, Spurr LF, Rouhani SJ, Chervin CS, Steinhardt G, Lager AM, Wanjari P, Malik R, Connell PP, Chmura SJ, Juloori A, et al. CDKN2A loss-of-function predicts immunotherapy resistance in non-small cell lung cancer. Sci Rep 2021;11:20059.

50. Ozenne P, Eymin B, Brambilla E, Gazzeri S. The ARF tumor suppressor: structure, functions and status in cancer. Int J Cancer 2010;127:2239–47.

51. Han G, Yang G, Hao D, Lu Y, Thein K, Simpson BS, Chen J, Sun R, Alhalabi O, Wang R, Dang M, Dai E, et al. 9p21 loss confers a cold tumor immune microenvironment and primary resistance to immune checkpoint therapy. Nat Commun 2021;12:5606.

52. Givechian KB, Garner C, Benz S, Song B, Rabizadeh S, Soon-Shiong P. An immunogenic NSCLC microenvironment is associated with favorable survival in lung adenocarcinoma. Oncotarget 2019;10:1840–9.

53. Binnewies M, Roberts EW, Kersten K, Chan V, Fearon DF, Merad M, Coussens LM, Gabrilovich DI, Ostrand-Rosenberg S, Hedrick CC, Vonderheide RH, Pittet MJ, et al. Understanding the tumor immune microenvironment (TIME) for effective therapy. Nat Med 2018;24:541–50.

54. Parra ER, Zhang J, Duose DY, Gonzalez-Kozlova E, Redman MW, Chen H, Manyam GC, Kumar G, Zhang J, Song X, Lazcano R, Marques-Piubelli ML, et al. Multi-omics Analysis Reveals Immune Features Associated with Immunotherapy Benefit in Patients with Squamous Cell Lung Cancer from Phase III Lung-MAP S1400I Trial. Clin Cancer Res 2024;30:1655–68.

55. William WN, Zhao X, Bianchi JJ, Lin HY, Cheng P, Lee JJ, Carter H, Alexandrov LB, Abraham JP, Spetzler DB, Dubinett SM, Cleveland DW, et al. Immune evasion in HPV-head and neck precancer-cancer transition is driven by an aneuploid switch involving chromosome 9p loss. Proc Natl Acad Sci U S A 2021;118.

