## Supplementary Info for "Gene signatures characterizing driver mutations in lung squamous carcinoma are predictive of the progression of pre-cancer lesions"

### Supplementary Materials

**Supplementary Table 1**: Frequent amplificated genes from similar chromosomal location

| Combined signature | Gene | Chr | Band | TCGA Sample Size |
|---|---|---|---|---|
| | ARNT | 1 | q21.3 | 56 |
| | TPM3 | 1 | q21.3 | 51 |
| | MUC1 | 1 | q22 | 53 |
| | FCGR2B | 1 | q23.3 | 55 |
| | PBX1 | 1 | q23.3 | 53 |
| chr1q21:23 | SDHC | 1 | q23.3 | 54 |
| | XPO1 | 2 | p15 | 62 |
| chr2p15:16 | REL | 2 | p16.1 | 66 |
| | TFG | 3 | q12.2 | 96 |
| chr3q12:13 | CBLB | 3 | q13.11 | 101 |
| | GATA2 | 3 | q21.3 | 112 |
| chr3q21:22 | FOXL2 | 3 | q22.3 | 139 |
| | WWTR1 | 3 | q25.1 | 203 |
| | GMPS | 3 | q25.31 | 214 |
| chr3q25 | MLF 1 | 3 | q25.32 | 227 |
| | PIK3CA | 3 | q26.32 | 295 |
| | SOX2 | 3 | q26.33 | 304 |
| | ETV5 | 3 | q27.2 | 283 |
| | EIF4A2 | 3 | q27.3 | 278 |
| | LPP | 3 | q27.3 | 273 |
| chr3q26:27 | BCL6 | 3 | q27.3 | 275 |
| | LIFR | 5 | p13.1 | 155 |
| chr5p13 | IL7R | 5 | p13.2 | 159 |
| | EXT1 | 8 | q24.11 | 71 |
| | RAD21 | 8 | q24.11 | 72 |
| | COX6C | 8 | q22.2 | 59 |
| | MYC | 8 | q24.21 | 96 |
| chr8q22:24 | NDRG1 | 8 | q24.22 | 77 |
| | WHSC1L1 | 8 | p11.23 | 108 |
| chr8p11 | FGFR1 | 8 | p11.23 | 103 |
| | ZNF384 | 12 | p13.31 | 52 |
| chr12p13 | CCND2 | 12 | p13.32 | 53 |

This table presents the unified amplification events identified by merging frequently amplified cancer genes, named based on their chromosomal locations in the genome. The following gene groups, corresponding to regions of frequent amplification, were defined: chr1q21:23, chr2p15:16, chr3q12:13, chr3q21:22, chr3q25, chr3q26:27, chr5p13, chr8q22:24, chr8p11, and chr12p13.

**Supplementary Table 2**: Prognostic validation on significant signatures in GSE157011

| Signature | survreg.pval | univariable HR |
|---:|---:|---:|
| MLL3-MUT | 0.005 | 1.23 |
| PIK3CA-MUT | 0.001 | 1.307 |
| TERT-AMP | 0.001 | 1.29 |
| chr5p13-AMP | 0.003 | 1.281 |
| chr8p11-AMP | 0.034 | 1.184 |
| chr8q22:24-AMP | 0.009 | 1.247 |
| CCND1-AMP | 0.007 | 1.274 |
| EGFR-AMP | 0.023 | 1.184 |
| ASXL1-AMP | 0.009 | 1.228 |
| HEY1-AMP | 0.00018652 | 1.356 |
| HOOK3-AMP | 0.004 | 1.249 |
| CDKN2A-DEL | 0.02 | 1.214 |
| FOXP1-DEL | 0.011 | 1.224 |
| PCM1-DEL | 0.039 | 1.162 |

Univariable Cox regression analysis in GSE157011 identified 14 signatures significantly associated with patient prognosis. The table lists the signature name, survival regression p-value (survreg.pval), and the univariable hazard ratio (univariable HR). All of these signatures were negatively correlated with survival (HR > 1), suggesting that the presence of these aberrations, or the deregulation of related pathways, detrimentally impacts patient prognosis.

| Signature | survreg.pval | multivariable HR |
|---:|---:|---:|
| MLL3-MUT | 0.033 | 1.178 |
| PIK3CA-MUT | 0.017 | 1.223 |
| TP53-MUT | 0.049 | 1.159 |
| TERT-AMP | 0.017 | 1.220 |
| IL7R-AMP | 0.026 | 1.214 |
| CCND1-AMP | 0.018 | 1.241 |
| EGFR-AMP | 0.019 | 1.210 |
| HEY1-AMP | 0.029 | 1.208 |

Multivariable Cox regression analysis in GSE157011 identified 8 signatures significantly associated with patient prognosis after adjusting for age, sex and stage. The table lists the signature name, survival regression p-value (survreg.pval), and the univariable hazard ratio (univariable HR). All of these signatures were negatively correlated with survival (HR > 1), suggesting that the presence of these aberrations, or the deregulation of related pathways, detrimentally impacts patient prognosis.

**Supplementary Table 3:** Correlation between signatures and immune regulatory genes

Supplementary Tables 3 represent the correlation between gene signatures and immune stimulators. Gene signatures include CDKN2A-DEL, CDKN2A-MUT, KRAS-AMP, SOX2-AMP, and TP53-MUT. Each row represents a developmental stage, and each column represents different immune stimulators. Correlation coefficients are visualized using a color gradient, where red indicates a positive correlation and blue indicates a negative correlation. This visualization highlights the dynamic interplay between gene expression signatures and immune regulation across developmental stages in LUSC.

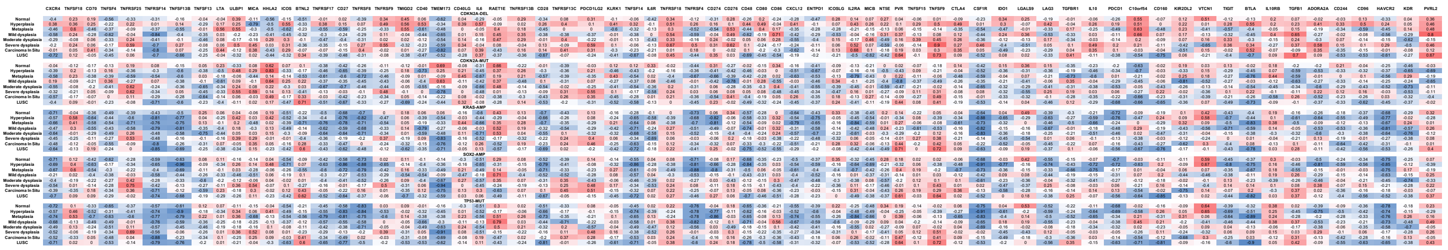

**Supplementary Table 4:** Functional annotation for driver genes

This table provides functional annotations for full set of 56 driver genes identified in this study. Each row corresponds to a specific gene, with the following columns: gene name, chromosomal location and a brief annotation of gene's role in oncogenic pathways, chromatin remodeling, transcriptional regulation, and cellular signaling.

**Supplementary Table 5**: Up-Regulated Genes Across Signatures
This table includes the top 50 genes with the highest absolute weights for each up-regulated gene expression signature. The columns represent distinct gene expression signatures, while the rows list the genes ranked by their weights. These genes are the most influential in defining each up-regulated signature.

**Supplementary Table 6:** Down-Regulated Genes Across Signatures
This table presents the top 50 genes with the highest absolute weights for each down-regulated gene expression signature. Similar to Supplementary Table 6, the columns represent different signatures, and the rows list the genes ranked by their weights, highlighting the key contributors to each down-regulated signature.

**Supplementary Table 7:** Co-deletion Frequency of CDKN2A and CD274 in TCGA-LUSC Samples

| | WT | CD274-DEL (36) |
|---|---|---|
| WT | 358(71.46%) | 3(0.6%) |
| CDKN2A-DEL(140) | 107(21.36%) | 33 (6.59)% |

This table summarizes deletion patterns in 501 TCGA-LUSC samples. CD274 deletion was rare and mostly co-occurred with CDKN2A deletion, suggesting it is part of broader 9p21.3 loss. In contrast, CDKN2A deletion frequently occurred independently.

Not displayed here due to overlarge size of the table.  Please see the file named “s4_56_GENE_ lusc.xlsx”, “S5_dn-regulated-gene.csv”, “S6_dn-regulated-gene.csv”

**Supplementary Figure 1**

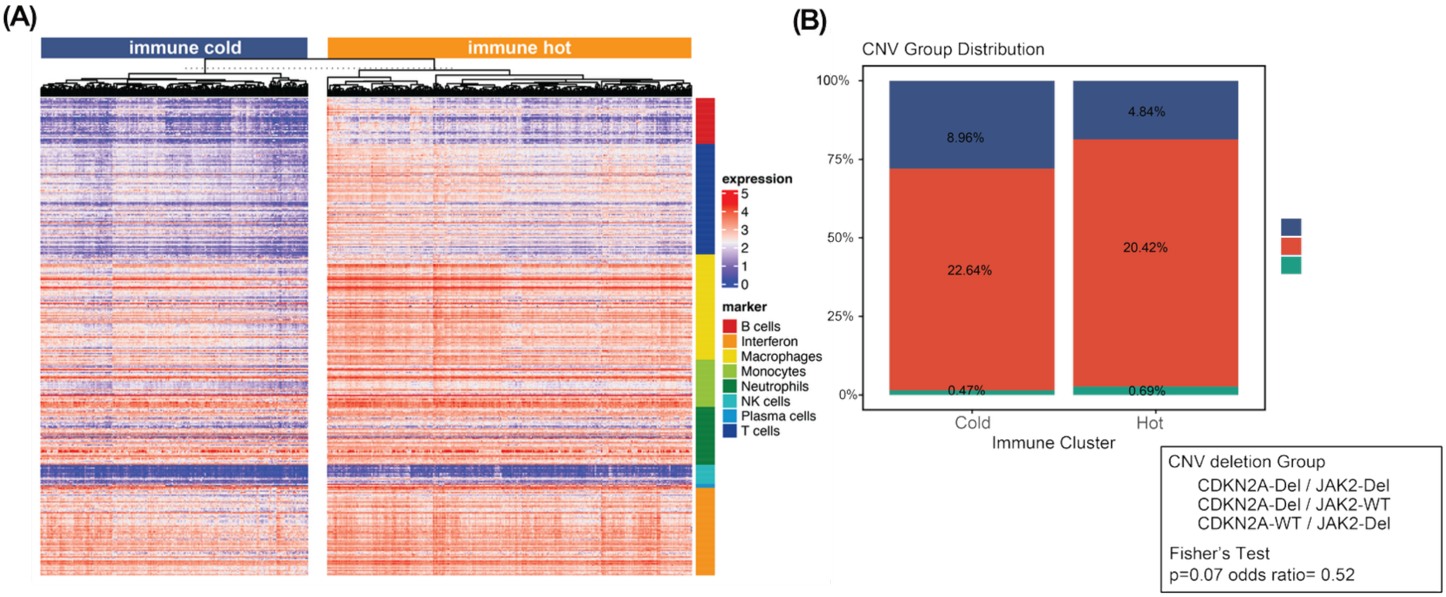


(A) In TCGA-LUSC data, heatmap showing samples categorized into the immune hot and immune cold groups, with hierarchical clustering of marker gene expression including B cells, interferon, macrophages, monocytes, NK cells, plasma cells, and T-cells. The rightmost color bar represents the groups for each marker gene group.

(B) Co-loss of CDKN2A and JAK2 deletions observed in 8.96% of immune cold samples versus 4.84% of immune hot samples. Patients are stratified by CDKN2A and JAK2 deletion status.

**Supplementary Note 1**

Define gene-signatures for driver genomic aberrations

We define gene signatures for each of these 34 driver genomic aberrations based on the LUSC transcriptomic data from TCGA. First, we perform a univariable regression model for each driver events to identify the informative genes, as in Eq.1:

(Eq. 1)

$$Y \sim \alpha + \sum_{i=1}^{m} \beta_i X_i$$

We encode a binary indicator variable for each genomic feature in each sample ($X_i = 1$: presence of mutation; $X_i = 0$: absence of mutation); Y is the $i_{th}$ gene expression; m is the total number of driver genomic aberrations. From the regression summary, we obtain the coefficients matrix $\beta_{i\times m}$ and the p-values matrix $P_{g\times m}$ for all of g genes and m events. Informative genes are those significant entries in $P_{g\times m}$ matrix.

Next, we define weight profiles consisting of all genes. We construct a pair of weighted profiles including $w^{up} = \left(w_1^{up}, w_2^{up}, \dots, w_g^{up}\right)$ and $w^{dn} = \left(w_1^{dn}, w_2^{dn}, \dots, w_g^{dn}\right)$ respectively for up-regulated and down-regulated genes. For a particular driver event with coefficient $\beta_k$ and p-value $P_k$, we define the weight by the following

(Eq. 2)

$$\begin{cases} \beta_k > 0 \ = \ -log_{10}\ (P_k) \\ \beta_k < 0 = log_{10}\ (P_k) \end{cases}$$

To avoid extreme values, we trim all weights greater than 10 to 10.

Calculate sample-specific gene signature score based on gene expression profiles

We use the previously developed computational algorithm BASE to calculate rank-based similarity score between TCGA-LUSC derived gene expression score and gene expression profile from independent validation datasets.[30] We follow these procedures: 1) Given an expression profile, we sort genes in the descending order as $e\ = (e_1, e_2, \dots, e_n)$, n is the total number of genes. 2) We validate whether genes with high weights shows skewed distribution and quantify the distribution difference via cumulative foreground function $f(i) = \frac{\sum_{k=1}^{i}|e_k w_k|}{\sum_{k=1}^{n}|e_k w_k|}, 1 \le i \le n$ and background function $b(i) = \frac{\sum_{k=1}^{i}|e_k(1-w_k)|}{\sum_{k=1}^{n}|e_k(1-w_k)|}, 1 \le i \le n$. 3) BASE algorithm then calculates maximum deviations between $f(i)$ and $b(i)$ against the permutated null distribution, resulting in $S^{up}$ and $S^{dn}$, respectively for $w^{up}$ and $w^{dn}$. The top genes are saved in S5 and S6 for 50 up-regulated genes and 50 down-regulated genes with max weights for each gene signature. The final gene-signature score is defined as:

(Eq. 4)

$$S\ =\ S^{up} - S^{dn}$$

where for each gene-signature, we determine the final score with the subtraction of the enrichment score of up-regulated models and down-regulated models for $g$. Extreme signature score ($S$ > 4 or $S < -4$) are set to 4, and -4 respectively.

**Supplementary Note 2**

Diagrammatic illustration to rationalize the use of gene expression signatures to characterize the oncogenic pathway de-regulation through diverse mechanisms (including primary and alternative genomic alterations) during tumorigenesis. The inactivation of the P53 pathway is selected as an example.

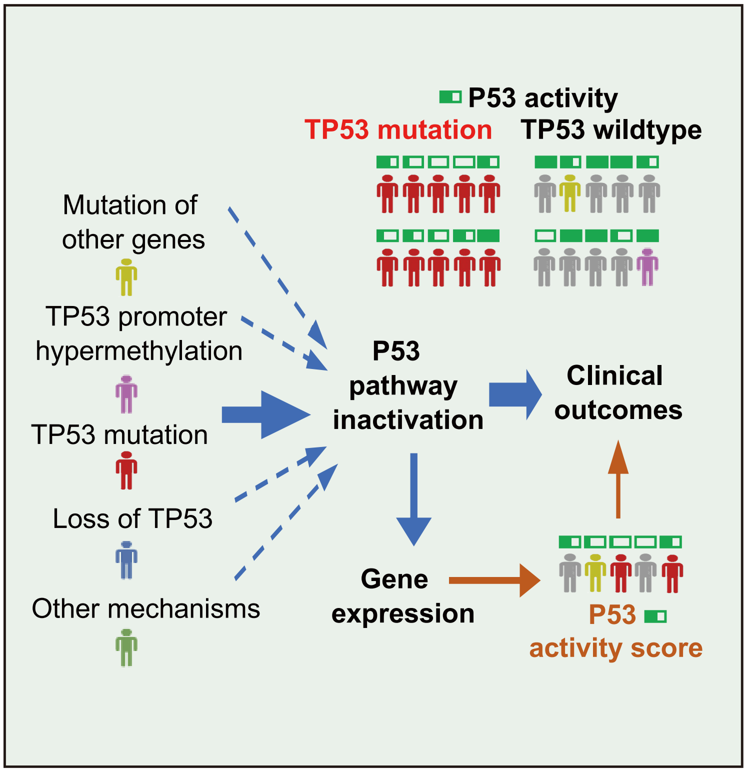
